# Relying on a declarative as opposed to a procedural learning strategy predisposes to the development of instrumental habits in female but not in male rats

**DOI:** 10.64898/2026.11.23.768427

**Authors:** Joseph S. Innes, David Belin

## Abstract

Across species, individuals differ in their reliance on declarative versus procedural long-term memory systems for exploring and controlling the environment, mapping respectively onto allocentric place versus egocentric response learning strategies in a spatial navigation task. Several factors motivate the widely held consideration that procedural spatial learning strategies are generalisable to stimulus-response habits in the domain of instrumental conditioning, and *vice versa*; however, this hypothesis has received little empirical testing. Here, in a prospective longitudinal study of 72 rats (36 females), each animal was characterised for its place/response learning tendency using a dual-solution T-maze task and a “direct opposition” test. Rats were then instrumentally conditioned to lever-press for food under a variable interval (VI) 60s schedule of reinforcement for five (*Experiment 1*) or 20 (*Experiment 2*) daily sessions, and the associative nature of their responding was assessed via sensitivity to offline reinforcer devaluation by pre-feeding to satiety. Against our hypothesis, response learners were less prone than place learners to rely on stimulus-response habits, an effect confined to female rats. Taken together, these findings reveal a complex, sex-dependent relationship between declarative and procedural memory systems and the two-fold associative basis of instrumental behaviour.

## Introduction

Convergent evidence from neuropsychological studies of human patients and controlled lesion studies in non-human animal models supports the current view that different types of learning and memory are subserved by dissociable neural systems operating in relative independence of each other (Cohen and Eichenbaum 1993; Milner et al. 1968; Scoville and Milner 1957; Squire 2004; White and McDonald 2002). A distinction paradigmatically made in this respect is that between declarative and procedural memory, demarcating the kind of memory the representational content of which can be separated from performance (“knowing that”) from that whose content cannot be so separated (“knowing how”), respectively (Cohen and Eichenbaum 1993; Cohen and Squire 1980; Squire 1986). While the neural substrate of declarative memory has been localised to the medial temporal lobe, including the hippocampus, and midline diencephalon (Aggleton and Brown 1999; Squire and Zola-Morgan 1991); that of procedural memory is instead a distributed network involving the striatum, the amygdala, the cerebellum and the neocortex (Bechara et al. 1995; Carlesimo et al. 1994; Daum et al. 1993; Fernandez-Ruiz et al. 2001; Kim and Thompson 1997; Knowlton et al. 1996; Phillips and LeDoux 1992; Squire 2004).

The distinction between declarative and procedural memory can be observed in the domain of spatial navigation: that is, humans and non-human animals can navigate towards goals using declarative and procedural learning strategies (Cohen and Eichenbaum 1993). The former requires the derivation of spatial parameters in an “allocentric” frame of reference that is not influenced by the animal’s own position and orientation. In a place learning strategy, animals learn the location of the goal in a coordinate system constructed from the geometry and cues in the navigational environment (O’Keefe and Nadel 1978; Packard and McGaugh 1996; Tolman et al. 1946). In contrast, for procedural learning strategies spatial parameters are derived in an “egocentric” frame of reference modulated by the animal’s position and orientation. In one procedural strategy, known as response learning, the animal learns to align or re-align its body axis (e.g., “turn left”) in response to specific cues it encounters along its path (O’Keefe and Nadel 1978; Packard and McGaugh 1996; Tolman et al. 1946).

Place and response learning strategies are subserved by dissociable neural systems, namely the dorsal hippocampus and dorsolateral striatum (DLS), respectively (Chang and Gold 2003, 2004; Morris et al. 1982; O’Keefe et al. 1975; Packard et al. 1989; Packard and McGaugh 1996).

In some “dual-solution” navigation tasks, both declarative and procedural strategies can provide the solution (Blodgett and McCutchan 1948; McDonald and White 1994). Here, the reliance on one as opposed to the other learning strategy to solve the task can be assessed in well-trained animals using a modified trial designed to exploit the behavioural distinction that “place learners” but not “response learners” will navigate towards the location of the goal with an efficient path when started from a novel location (Blodgett and McCutchan 1948; Packard and McGaugh 1996; Restle 1957).

The reliance on place as opposed to response navigation is controlled by various situational factors (Packard and Goodman 2013). For example, with extensive navigational practice, animals initially relying on a place strategy tend to switch to rely on a response strategy (Hicks 1964; Iaria et al. 2003; Mackintosh 1965; Packard and McGaugh 1996). However, the reliance on place versus response learning to solve a navigation task is further influenced by genetic polymorphisms (Konishi et al. 2016) and allied to behavioural traits shown to have strong test-retest consistency (Adamec and Shallow 2000; Grissom et al. 2012; Hawley et al. 2011). As such, in well-controlled experiments, we can probably infer from the reliance on place versus response learning an individually consistent phenotype, *place/response learning tendency*.

Similarly, instrumental learning depends on two processes, namely goal-directed actions and stimulus-response (S-R) habits, which are underpinned by distinct associative structures and dissociable neural substrates (Balleine and O’Doherty 2010; Dickinson 1985). Instrumental behaviour, that is, behaviour acquired and maintained under the control of its consequences (Dickinson 1994), can be said to be goal-directed when it is under the integrated control of *(i)* the present causal relationship between behaviour and outcome and *(ii)* the incentive value of that outcome (Dickinson 1985; Dickinson and Balleine 1994; de Wit and Dickinson 2009). It is generally considered to have an action-outcome (A-O) associative structure and to enable adaptive, flexile behaviour (Dickinson 1985; Dickinson and Balleine 1994; de Wit and Dickinson 2009). In contrast, S-R habits are instrumental behaviours acquired and maintained under the control of an association between antecedent setting and response, an S-R association, strengthened by contiguous delivery of a valued outcome (Dickinson 1985; Thorndike 1898, 1911). Habits are therefore bound to settings and less cognitively taxing, but also less flexible than goal-directed behaviours (Wood and Rünger 2016; Thrailkill and Bouton 2015). Thus, when the instrumental outcome is devalued offline, for instance by pre-feeding it to satiety (Colwill and Rescorla 1985), with all else equal, only animals relying on goal-directed actions will reduce their response rate for that outcome when assayed under extinction conditions, under which conditions animals cannot procure, and hence experience, the devalued outcome contiguously to the response.

As occurs with regard to place and response learning strategies in dual-solution navigation tasks (see above), with extensive training animals relying on goal-directed control over instrumental behaviour switch to rely on instrumental habits, at least under schedules of reinforcement with a low response-outcome contingency (DeRusso 2010; Dickinson et al. 1983; Garr et al. 2021; Tricomi et al. 2009; but see in humans de Wit et al. 2018; Pool et al. 2022). This is paralleled by a shift in the neural locus of control over instrumental behaviour across parallel dissociable memory systems, including a transition from the engagement of dopamine-dependent mechanisms in the posterior dorsomedial striatum (pDMS) to those in the anterior dorsolateral striatum (aDLS; Corbit and Janak 2010; Yin et al. 2004, 2006, 2005; Faure et al. 2005; Murray et al. 2012; Corbit et al. 2012).

Similarities in the influence of factors such as training and underlying neural substrates motivate the hypothesis that response learning and S-R habits, on one hand, and even place learning and goal-directed actions, on the other, are the same behavioural constructs applied to different domains of spatial navigation and instrumental conditioning (Kosaki et al. 2018); or, at least, that they are underpinned by the same multiple memory systems architecture (Chersi and Burgess 2015; Geerts et al. 2020). Indeed, response navigation is routinely used to study S-R habits (e.g., Ferragud et al. 2010; Packard 1999; Packard and McGaugh 1996; Smith et al. 2012; Smith and Graybiel 2013). This hypothesis, however, has rarely been directed tested (but see Kosaki et al. 2018; De Leonibus et al. 2011; Sage and Knowlton 2000; Vikbladh et al. 2019).

One approach has assessed whether response and place navigation meet the behavioural criteria for S-R habits and goal-directed actions. Some studies have suggested that well-trained spatial performance under a response as opposed to a place strategy is relatively resistant to offline changes in the incentive value of the outcome, motivating the conclusion that response and place learning “each conform to one of the definitions of S–R habit and goal-directed action, respectively” (Kosaki et al. 2018; see also Sage and Knowlton 2000). However, this particular inference relies on the assumption that spatial navigation under place and response learning is an instrumental behaviour, which has not yet been demonstrated (Dickinson 1994).

An alternative approach, which is to be employed in the present study, tests whether the individual tendency to rely on response as opposed to place learning in spatial navigation predisposes to an increased tendency to rely on instrumental habits as opposed to goal-directed actions in an instrumental context. Indeed, in humans, the reliance on procedural cue-driven versus declarative boundary-driven strategies in a virtual reality navigation task has been shown to predict the reliance on goal-directed model-based versus habitual model-free strategies in a two-step probabilistic decision-making task (Vikbladh et al. 2019). However, this result may be explained by the fact that this specific goal-directed learning process required the engagement of the declarative memory system to chain multiple behaviours across steps (Miller et al. 2017), as opposed to any correspondence between multiple memory systems subserving distinct spatial and instrumental learning strategies. This will not be the case for tasks more conventionally used to assess the reliance on goal-directed as opposed to habitual behaviour, such as those employing the free-operant design (Ferster 1953).

Therefore, in the present series of longitudinal prospective experiments, two cohorts of male and female outbred rats were screened for their place/response learning tendency before being trained instrumentally to respond for an ingestive reinforcer, after which the goal-directed versus habitual nature of instrumental behaviour was assessed. Chiefly, rats were trained in a T-maze spatial discrimination task that can be solved using place and response learning strategies, and the strategy on which animals relied to solve the discrimination was assayed after training in a “direct opposition” test in which the maze was rotated by 180° (Blodgett and McCutchan 1948; Packard and McGaugh 1996; Restle 1957). Subsequently, place and response learners were given limited (*Experiment 1*) or extended (*Experiment 2*) training in a free-operant instrumental task under a schedule of reinforcement that promotes the development of S-R habits (Dickinson et al. 1983). The goal-directed versus habitual nature of instrumental behaviour was assayed by measuring under extinction conditions its sensitivity to the offline devaluation of the reinforcer by pre-feeding to satiety (Colwill and Rescorla 1985; Gourley et al. 2010).

## Results

The experiments of this study are schematised in **Figure 1**.

**Figure 1:**
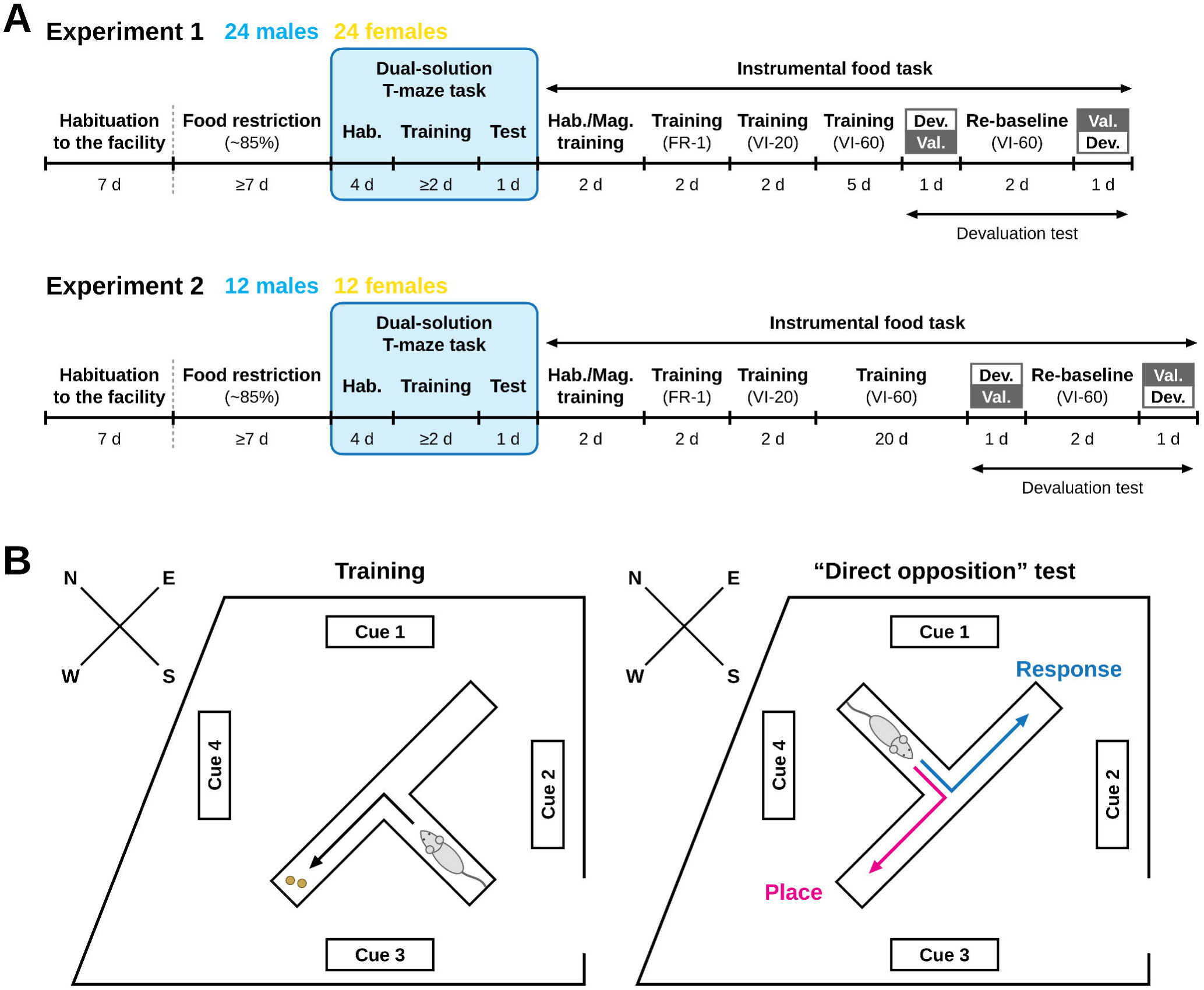
Schematic summary of the experiments. **A)** After a seven-day period of habituation to the animal facility, a total of 72 rats (36 females) were food-restricted to reduce their body weight to ∼85% of their predicted free-feeding values. Subsequently, hungry rats were trained in a spatial discrimination task in a T-maze, being classified as place learners or response learners depending on their behaviour in a post-training “direct opposition” test (see *B*). Afterwards, rats were trained to respond instrumentally for food by pressing a lever under a variable interval (VI) 60s schedule of reinforcement for either five (*Experiment 1*) or 20 (*Experiment 2*) daily sessions. In a post-training, within-subjects devaluation test, rats underwent two five-minute tests conducted under extinction conditions and separated by two daily sessions of VI 60s training to re-baseline instrumental performance. Before one of the two tests (counterbalanced), the reinforcer was devalued by pre-feeding to satiety. Before the remaining test, rats received no intervention (i.e., the valued condition). **B)** In the dual-solution T-maze task, hungry rats were trained to retrieve food from one of the two lateral arms of the T-maze. This spatial discrimination could be solved using a place learning strategy, where rats learn the location of the goal in an allocentric frame of reference built from the geometry of and extra-maze cues within the testing room, or a response learning strategy, where rats learn to execute a turning response in response to specific intra-maze cues along their path to the choice point. To determine the strategy on which each subject relied to solve the spatial discrimination, a “direct opposition” test was conducted 24 hours after learning criterion was achieved in training. This trial, conducted under extinction conditions, was performed with the maze rotated by 180°. “Place learners” and “response learners” are expected to select opposite lateral arms of the maze under these conditions. Dev. = devalued; FR = fixed ratio; Hab. = habituation; Mag. Training = magazine training; Val. = valued.

Male and female rats acquired the T-maze spatial discrimination to the level of accuracy specified by the learning criterion (i.e., five consecutive correct trials) within 15.66 ± 11.12 (*Experiment 1*) and 20.40 ± 7.65 (*Experiment 2*) trials. Males and females required the same number of trials to attain the learning criterion (Sex (*Experiment 1*): F_1,37_ = 1.02, p = 0.32, η_p_^2^ = 0.027; Sex (*Experiment 2*): F_1,16_ < 1). When the navigation strategy, place or response, on which rats relied to solve the dual-solution T-maze task was determined in a “direct opposition” test (see *Methods and materials*), 13 out of 41 (*Experiment 1*) and nine out of 20 (*Experiment 2*) subjects displayed a reliance on a place learning strategy and were classified as “place learners”, while the remainder relied on a response strategy and were classified as “response learners” (**Figure 2A**, **Figure 3A**). Although response learners were slightly greater represented in males than in females in *Experiment 1* (**Figure 2A**), statistical analysis suggested that the distribution of place and response learners did not depend on sex in either experiment (**Figure 2A**, **Figure 3A**; *Experiment 1*: χ^2^(1) = 1.86, p = 0.17, OR = 2.53; *Experiment 2*: p = 1.00, OR = 1.04). Place and response learners required the same number of trials to attain the learning criterion in the dual-solution T-maze task (**Figure 2B**, **Figure 3B**; for all F, F < 1); this suggests that individual differences in the reliance on place as opposed to response learning cannot simply be explained by variation in the quantity of training received.

**Figure 2:**
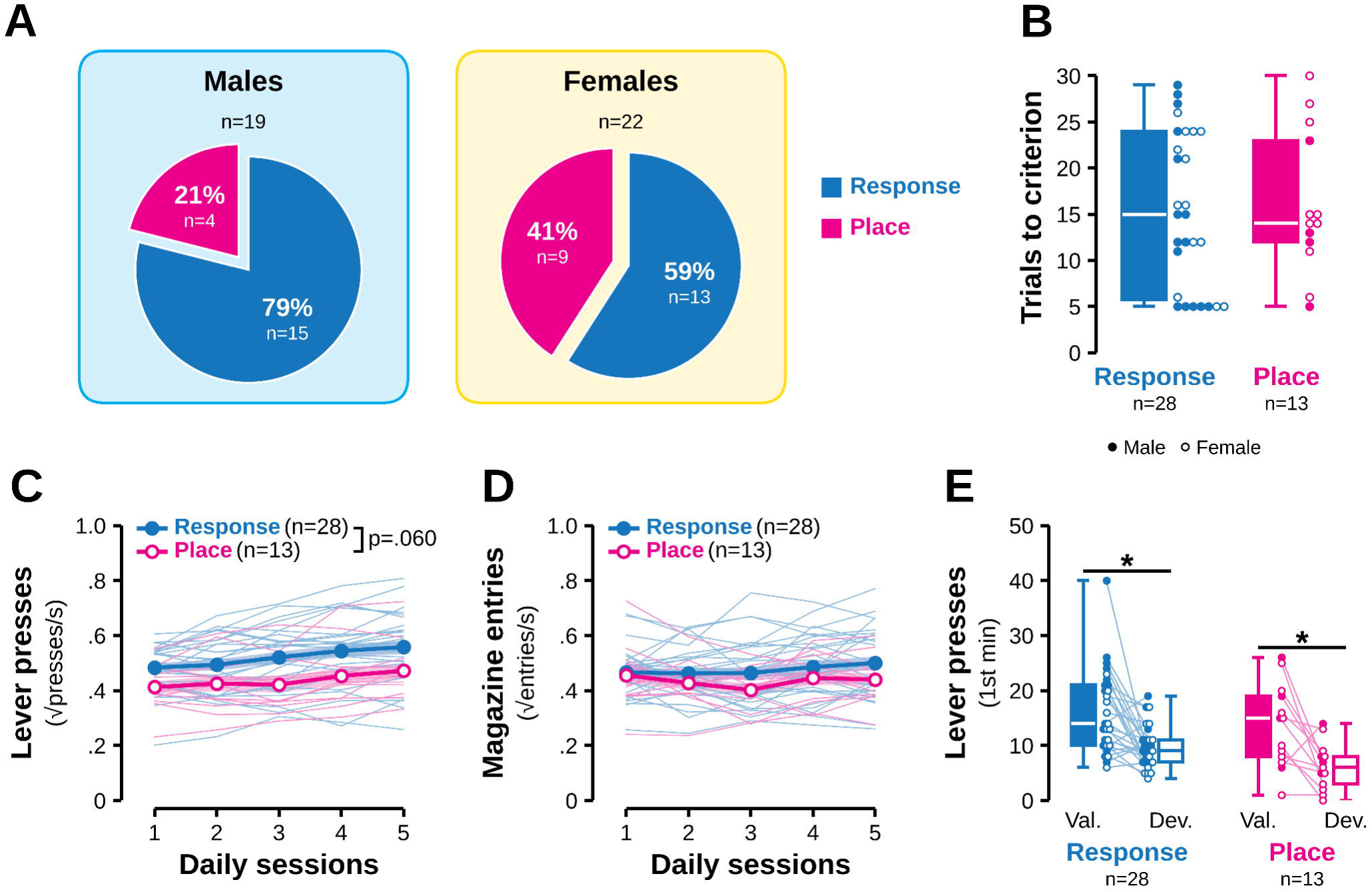
Place and response learners identified in a cohort of male and female rats differ marginally in their instrumental response rate under a variable interval schedule of reinforcement but are equally sensitive to reinforcer devaluation after limited training. **A)** Out of the 41 rats that attained criterion level of discrimination accuracy in the dual-solution T-maze task, 13 were identified as place learners and 28 as response learners in a post-training “direct opposition” test performed under extinction conditions and with the T-maze rotated by 180°. Females were slightly more likely to be place learners than males, but this differential representation was not statistically significant (χ^2^(1) = 1.86, p = 0.17, odds ratio (OR) = 2.53). **B)** Place and response learners acquired the T-maze spatial discrimination at the same rate, taking a similar number of trials to achieve the learning criterion (for all F, F < 1). **C)** Response learners pressed the lever slightly more frequently than place learners as subjects were instrumentally conditioned to respond for food under a variable interval schedule of reinforcement (Navigation Strategy: F_1,37_ = 3.77, p = 0.060, η ^2^= 0.092; Sex x Navigation Strategy: F_1,37_ < 1; Navigation Strategy x Day: F_2.29,84.73_ < 1; Sex x Navigation Strategy x Day: F_2.29,84.73_ = 1.04, p = 0.36, η_p_^2^= 0.027). **D)** Non-instrumental behaviours, such as magazine entries, were equally frequent between groups (Navigation Strategy: F_1,37_ < 1; Sex x Navigation Strategy: F_1,37_ < 1; Day: F_2.70,99.72_ = 1.84, p = 0.15, η_p_^2^ = 0.047; Navigation Strategy x Day: F_2.70,99.72_ = 1.30, p = 0.28, η_p_^2^ = 0.034; Sex x Navigation Strategy x Day: F_2.70,99.72_ = 1.40, p = 0.25, η ^2^ = 0.036). **E)** In a post-training devaluation test, the pre-feeding to satiety of the reinforcer (the devalued condition; Dev.) lowered the lever pressing rate during the first minute of a five-minute test conducted under extinction conditions as compared to no pre-feeding (the valued condition; Val.; Condition (min1): F_1,37_ = 21.99, p < 0.001, η_p_^2^ = 0.37). The effect of reinforcer devaluation on the instrumental response rate did not differ between rats subdivided according to their place/response learner identity (Navigation Strategy x Condition (min1): F_1,37_ = 2.06, p = 0.16, η_p_ ^2^ = 0.053; Sex x Navigation Strategy x Condition (min1): F_1,37_ < 1).

**Figure 3:**
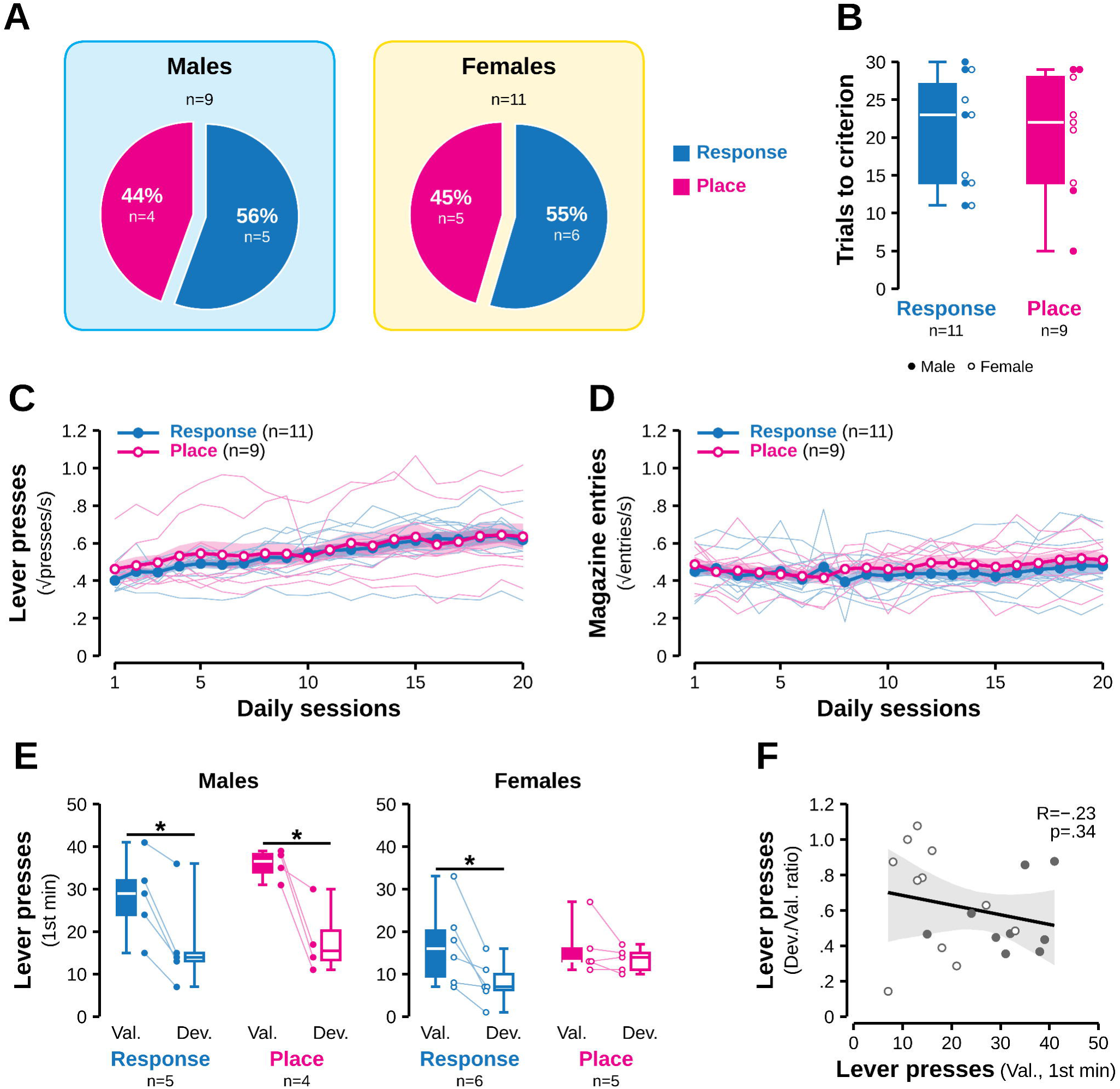
The reliance on place learning specifically confers a resistance of instrumental responding to reinforcer devaluation after extended training in females but not in males. **A)** Out of the 20 rats that attained criterion level of discrimination accuracy in the dual-solution T-maze task, nine were identified as place learners and 11 as response learners in the “direct opposition” test performed 24 hours after training under extinction conditions and with the T-maze rotated by 180°. Males and females were equally likely to be place or response learners (p = 1.00, odds ratio (OR) = 1.04). **B)** Place and response learners acquired the T-maze spatial discrimination at the same rate, taking a similar number of trials to achieve the learning criterion (for all F, F < 1). **C-D)** Place and response learners responded instrumentally at the same frequency throughout all 20 daily variable interval (VI) 60s sessions (Navigation Strategy: F_1,16_ < 1; Sex x Navigation Strategy: F_1,16_ = 2.14, p = 0.16, η_p_^2^ = 0.12; Navigation Strategy x Two-Day Block: F_2.72,43.49_ < 1; Sex x Navigation Strategy x Two-Day Block: F_2.72,43.49_ = 1.02, p = 0.39, η ^2^ = 0.060). They also entered the food magazine at an equal rate throughout this same training period (for all F, F < 1). **E)** In a post-training devaluation test, the pre-feeding to satiety of the reinforcer (the devalued condition; Dev.) reduced the rate of lever pressing during the first minute of a five-minute test conducted under extinction conditions compared to no pre-feeding (the valued condition; Val.; Condition (min1): F_1,16_ = 51.40, p < 0.001, η_p_^2^ = 0.76). Here, after extended training under a VI 60s schedule of reinforcement, the reduction in lever pressing after reinforcer devaluation as compared to after the control condition was less pronounced in place learners than in response learners, an effect confined to females (Navigation Strategy x Condition (min1): F_1,16_ = 1.06, p = 0.32, η ^2^ = 0.062; Sex x Navigation Strategy x Condition (min1): F = 5.15, p = 0.037, η_p_^2^ = 0.24). Indeed, whereas devaluation reduced the rate of instrumental responding in male place learners (p < 0.001), male response learners (p = 0.001), and female response learners (p = 0.003), there was no evidence that instrumental responding in place learners was sensitive to the same manipulation (p = 0.36). **F)** There was no dimensional relationship between the sensitivity of instrumental responding to reinforcer devaluation and the baseline response rate (i.e., that under the valued condition; *r*_19_ =-0.23, p = 0.34), suggesting that a floor effect cannot account for the present results.

Subjects classified as place and response learners were then trained instrumentally to respond by pressing a lever for 45 mg food pellets under a VI 60s schedule of reinforcement. Across a limited training period of five daily sessions (*Experiment 1*) or an extended training period of 20 daily sessions (*Experiment 2*), the instrumental response rate was either marginally greater in response learners than in place learners (*Experiment 1*; **Figure 2C**; Navigation Strategy: F_1,37_ = 3.77, p = 0.060, η_p_ ^2^ = 0.092; Sex x Navigation Strategy: F_1,37_ < 1; Navigation Strategy x Day: F_2.29,84.73_ < 1; Sex x Navigation Strategy x Day: F_2.29,84.73_ = 1.04, p = 0.36, η_p_^2^ = 0.027) or equal across groups (*Experiment 2*; **Figure 3C**; Navigation Strategy: F_1,16_ < 1; Sex x Navigation Strategy: F_1,16_ = 2.14, p = 0.16, η_p_ ^2^ = 0.12; Navigation Strategy x Two-Day Block: F_2.72,43.49_ < 1; Sex x Navigation Strategy x Two-Day Block: F_2.72,43.49_ = 1.02, p = 0.39, η_p_^2^ = 0.060). Behaviours that were not explicitly instrumentally conditioned, such as magazine entries, were executed at the same rate in place and response learners throughout both experiments (**Figure 2D**, **Figure 3D**; Navigation Strategy (*Experiment 1*): F_1,37_ < 1; Sex x Navigation Strategy (*Experiment 1*): F_1,37_ < 1; Day (*Experiment 1*): F_2.70,99.72_ = 1.84, p = 0.15, η_p_^2^ = 0.047; Navigation Strategy x Day (*Experiment 1*): F_2.70,99.72_ = 1.30, p = 0.28, η_p_^2^ = 0.034; Sex x Navigation Strategy x Day (*Experiment 1*): F_2.70,99.72_ = 1.40, p = 0.25, η_p_^2^ = 0.036; for all F (*Experiment 2*), F < 1).

Consistent with previous studies (Quinn et al. 2007), male subjects responded instrumentally for food rewards at a greater frequency than females throughout instrumental training under a VI 60s schedule of reinforcement (**Supplementary Figure 1**, **Supplementary Figure 2**).

After limited conditioning, instrumental responding was, as expected, markedly disrupted by the offline devaluation of the reinforcer by pre-feeding to satiety (**Figure 2E**; Condition (min1): F_1,37_ = 21.99, p < 0.001, η_p_^2^= 0.37). Here, reinforcer devaluation attenuated responding under extinction conditions equally in place and response learners (**Figure 2E**; Navigation Strategy x Condition (min1): F_1,37_ = 2.06, p = 0.16, η_p_^2^ = 0.053; Sex x Navigation Strategy x Condition (min1): F_1,37_ < 1). After an extended period of instrumental conditioning in *Experiment 2*, responding under extinction conditions was also disrupted by the pre-feeding to satiety of the instrumental outcome (**Figure 3E**; Condition (min1): F_1,16_ = 51.40, p < 0.001, η_p_^2^= 0.76). However, this time, place and response learners were differentially sensitive to reinforcer devaluation, an effect confined to females (**Figure 3E**; Navigation Strategy x Condition (min1): F_1,16_ = 1.06, p = 0.32, η_p_ ^2^ = 0.062; Sex x Navigation Strategy x Condition (min1): F_1,16_ = 5.15, p = 0.037, η_p_^2^ = 0.24). Whereas, in males, lever pressing was equally attenuated by devaluation across groups (place learners: p < 0.001; response learners: p = 0.001), instrumental responding was reduced in female response learners (p = 0.003) but not in their place learner counterparts (p = 0.36). This effect could not be explained by variability in the baseline instrumental response rate (e.g., due to a floor effect) as the sensitivity of lever pressing to devaluation, expressed as a ratio, was not dimensionally associated with the lever pressing rate during testing under the valued condition (**Figure 3F**; *r*_19_ =-0.23, p = 0.34).

Further analyses suggested that such group differences in devaluation sensitivity were specific to the instrumentally conditioned behaviour. A comparison of the free consumption of the reinforcer under devalued and valued conditions, measured in a 15-minute test immediately after the instrumental response rate was assessed, was taken as proxy of the efficacy of the pre-feeding to satiety-mediated devaluation procedure. Although such consumption was profoundly attenuated by reinforcer devaluation, this reduction in free outcome consumption was of the same magnitude in place and response learners (**Figure 4A**; Condition: F_1,16_ = 24.87, p < 0.001, η_p_^2^= 0.61; Navigation Strategy x Condition: F_1,16_ < 1; Sex x Navigation Strategy x Condition: F_1,16_ < 1). Similarly, non-instrumental behaviours, i.e., magazine entries, were reduced by devaluation regardless of place/response learner identity (**Figure 4B**; Condition (min1): F_1,16_ = 20.40, p < 0.001, η_p_^2^ = 0.56; Navigation Strategy x Condition (min1): F_1,16_ = 1.16, p = 0.30, η_p_^2^ = 0.067; Sex x Navigation Strategy x Condition (min1): F_1,16_ < 1).

**Figure 4:**
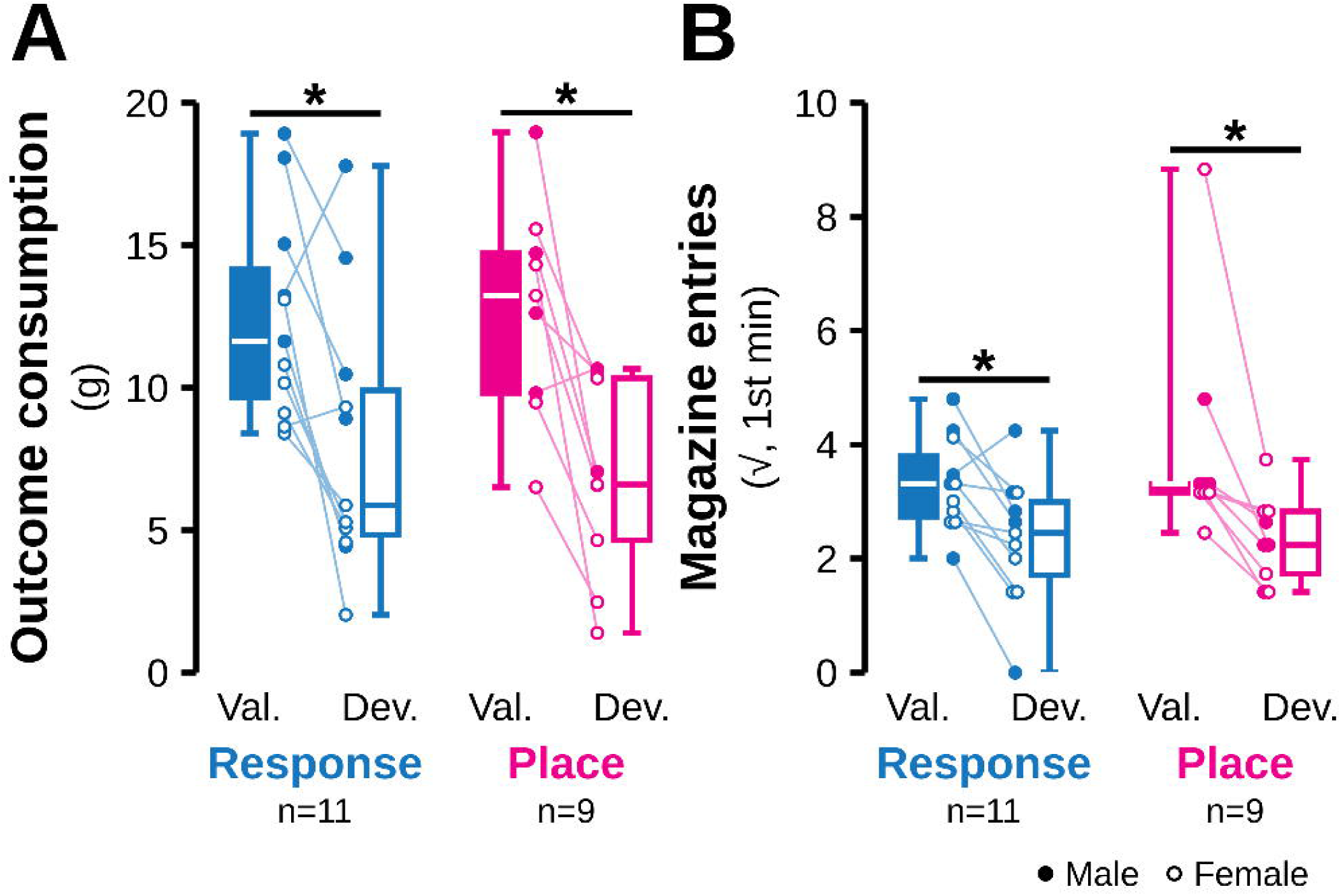
Pre-feeding to satiety of the reinforcer equally reduces the free consumption of the reinforcer and the rate of magazine entries in place and response learners. **A)** As expected, rats freely consumed a smaller quantity of the reinforcer after it was pre-fed to satiety (the devalued condition; Dev.) than after no intervention (the valued condition; Val.), with this being measured across a 15-minute assay starting immediately after the instrumental response rate was assessed in the devaluation test (Condition: F_1,16_ = 24.87, p < 0.001, η_p_^2^ = 0.61). The effect of reinforcer devaluation on the free consumption of the instrumental outcome did not differ between place learners and response learners: the two groups consumed equal quantities under devalued and valued conditions (Navigation Strategy: F_1,16_ < 1; Sex x Navigation Strategy: F_1,16_ = 1.58, p = 0.23, η ^2^ = 0.090; Navigation Strategy x Condition: F_1,16_ < 1; Sex x Navigation Strategy x Condition: F_1,16_ < 1). **B)** After 20 daily sessions of instrumental conditioning under a VI 60s schedule of reinforcement, non-instrumental behaviours such as magazine entries were sensitive to the devaluation of the instrumental outcome. That is, when compared to no intervention, the pre-feeding to satiety of the reinforcer reduced the rate of magazine entries during the first minute of a five-minute test conducted under extinction conditions (Condition (min1): F_1,16_ = 20.40, p < 0.001, η_p_^2^ = 0.56). This effect of devaluation was observed equally in place and response learners (Navigation Strategy x Condition (min1): F_1,16_ = 1.16, p = 0.30, η_p_^2^ = 0.067; Sex x Navigation Strategy x Condition (min1): F_1,16_ < 1).

In line with previous studies (Corbett et al. 2025; Mohammed et al. 2025; Quinn et al. 2007; Schoenberg et al. 2019, 2022), females were generally less sensitive to reinforcer devaluation in their instrumental responding than males after extended instrumental conditioning, although the pre-feeding to satiety of the reinforcer reduced the rate of lever pressing in both sexes (**Supplementary Figure 2**). In marked contrast, after limited conditioning, the sexes were equally sensitive to reinforcer devaluation in their lever pressing (**Supplementary Figure 1**).

## Discussion

The present study tested the hypothesis that rats tending to rely on a procedural response over a declarative place learning strategy for spatial navigation are more predisposed to develop S-R habitual control over instrumental behaviour. Using the sensitivity to the offline devaluation of the reinforcer after different histories of instrumental conditioning as a proxy for the reliance on goal-directed actions versus S-R habits of rats previously characterised as place or response learners in a T-maze task, the present results revealed that place learners were *more prone* than response learners to develop S-R habitual control after extended training, an effect that was restricted to female rats. This surprising finding went against our original hypothesis.

The present experiments used a dual-solution spatial discrimination task in the T-maze followed by a “direct opposition” test to divide cohorts of rats into those relying more on declarative spatial learning strategies, place learners, and those relying more on procedural strategies, response learners. While place and response learners were evenly distributed in female rats (**Figure 2A**, **Figure 3A**), consistent with previous studies using gonadally-intact animals (Korol et al. 2004), response learners were a numerical majority in males, particularly in *Experiment 1* (**Figure 2A**, **Figure 3A**). This finding, which we have replicated in our experimental set-up (J. S. Innes, unpublished findings), goes against most previous studies in which place learners predominate in male rats after a similar amount of training (e.g., Colombo et al. 2003; Hicks 1964; Mackintosh 1965; Packard 1999; Packard and McGaugh 1996).

That said, the individual reliance on declarative versus procedural spatial learning strategies is controlled by a range of situational factors that may vary from one navigational environment to the next (see *Introduction*). Of particular note, the large, near-square room in which the experiments took place and the placement of the T-maze within this room such that the discrimination axis ran along it’s diagonal (**Figure 1B**; see *Materials and methods*) may have discouraged the reliance on the environmental geometry instead of salient visual landmarks for allocentric spatial navigation. As male rats show a particular preference for allocentric navigation by geometry as opposed to landmarks (Williams et al. 1990), a greater fraction of such animals may have defaulted to response learning in this navigational environment.

The specific reinforcer devaluation protocol used in these experiments raises questions as to whether group differences in the reliance on S-R habits versus goal-directed actions can be validly inferred from those in the sensitivity of instrumental responding to devaluation elicited here. The logic of reinforcer devaluation is that S-R-mediated instrumental behaviour is not controlled by the current incentive value of the instrumental outcome and should not be affected by changes in such value insofar as the animal cannot subsequently produce the devalued outcome contiguously with the behaviour (Dickinson 1985; Dickinson and Balleine 1994; de Wit and Dickinson 2009). “Gold-standard” reinforcer devaluation protocols, i.e., conditioned taste aversion (CTA; Adams and Dickinson 1981; Garcia et al. 1955; Garcia and Koelling 1966) and sensory-specific satiety (Colwill and Rescorla 1985; Rolls 1986; Rolls et al. 1981), compare instrumental behaviour between two conditions across which only the incentive value of the reinforcer changes. The present reinforcer devaluation protocol also reduces such incentive value (Balleine and Dickinson 1998). However, since subjects were not satiated with a distinct food type under the valued condition (see *Methods and materials*), it also produces a general reduction in the primary motivational state of hunger that could disrupt instrumental behaviour (Dickinson et al. 1995). In principle, this could be achieved by four mechanisms.

1) A general reduction in locomotor activity (Campbell and Sheffield 1953; Wald and Jackson 1944).
2) A reduction in the rate of Pavlovian conditioned approach to the magazine, known as “goal-tracking” (Boakes 1977; Flagel et al. 2009). It could be envisaged that, during instrumental conditioning, goal-tracking behaviours facilitate instrumental responding indirectly by bringing animals into closer proximity with the *operandum*, i.e., the lever.
3) A reduction in the facilitatory effect of conditioned Pavlovian associations on the performance of an instrumentally conditioned behaviour, which is known as Pavlovian-to-instrumental transfer (PIT; Estes 1943; Lovibond 1983). PIT can be behaviourally and neurally dissociated into general and specific kinds (Corbit et al. 2007; Corbit and Balleine 2005). Under extensive instrumental conditioning in a free-operant task using a single *operandum*, general PIT appears to be dominant (Cartoni et al. 2016). General but not specific PIT is disrupted by downshifts in the primary motivational state related to the Pavlovian unconditioned stimulus (US; Corbit et al. 2007; Lingawi et al. 2022).
4) A change in “interoceptive context” generated by an alteration in primary motivational state. There is evidence that hunger states can operate similarly to “exteroceptive contexts”, for instance by mediating the “ABA” renewal of extinguished instrumental responding (Schepers and Bouton 2017; Thomas et al. 2023). Unlike that of Pavlovian behaviours (Bouton and King 1983; Bouton and Peck 1989), the performance of instrumentally conditioned behaviours is generally disrupted by switches in “exteroceptive” context (Bouton et al. 2011; Todd 2013), and this is amplified when such responses are under S-R associative control (Thrailkill and Bouton 2015). Contextual control over instrumental performance appears to be mediated predominantly by direct S-R (i.e., context-R) associations (Bouton and Todd 2014).

The potential involvement of at least one of these mechanisms in the present experiments is supported by the fact that even after 20 daily sessions of instrumental conditioning under lean VI schedules of reinforcement – sufficient to engender devaluation-resistant instrumental responding assessed using “gold-standard” assays (DeRusso 2010; Dickinson et al. 1983, 1995) – the majority of rats in Experiment 2 reduced their rate of instrumental responding after reinforcer devaluation compared to the baseline condition (**Figure 2E**). However, it is our contention that none of these mechanisms can adequately account for the place/response learning tendency-dependent differential sensitivity to devaluation demonstrated in this work.

The contribution of mechanisms *1)* and *2)* to our main findings is rendered unlikely by the result that behaviours not explicitly instrumentally conditioned, such as magazine entries, were equally disrupted by reinforcer devaluation in place and response learners (**Figure 4B**). The involvement of mechanism *3)* is ruled out by the fact that the differential devaluation-sensitivity of instrumental responding in female rats according to their place/response learning tendency was observed after extended but not after limited training (**Figure 2E**, **Figure 3E**). In the only study to investigate the contribution of general PIT to the effect of devaluation on instrumental responding, the size of such effect was equally pronounced after four and 20 daily sessions of instrumental conditioning under a VI 60s schedule of reinforcement (Jonkman et al. 2010).

Further, the design of the present experiments renders it unlikely that mechanism *4)* accounts in any significant way for the main findings of this study. Animals were food-deprived at least seven days prior to the start of instrumental conditioning (**Figure 1A**). Such pre-exposure should substantially reduce the associability of context through latent inhibition (Lubow 1973) and consequently the impact of “interoceptive context switch” on the instrumental response rate. This contrasts to Bouton and colleagues’ experiments, in which animals were food-deprived only on the days of instrumental conditioning (“A”) and renewal (“A”), rendering the associability of “interoceptive context” high (Schepers and Bouton 2017; Thomas et al. 2023).

As such, the results of this study probably reflect a differential sensitivity of instrumentally conditioned to an offline reduction in the incentive value of the reinforcer dependent on place/response learning tendency. This is further supported by the present finding that females are more devaluation-resistant in their instrumental responding than males after extensive conditioning (**Supplementary Figure 2**), in line with studies using “gold-standard” protocols for devaluation (Mohammed et al. 2025; Quinn et al. 2007; Schoenberg et al. 2019, 2022).

It would nonetheless reinforce our interpretation of the results of this study to show the same results using a “gold-standard” devaluation protocol. However, conditioning a taste aversion to an ingestive reward has profound, long-lasting effects on animals’ evaluation of that outcome (Garcia et al. 1955), and therefore CTA is a less amenable protocol for within-subjects testing of the habitual nature of instrumental behaviour, such as the longitudinal study of individual differences in the present experiments. As a result, sensory-specific satiety, which has short-lasting effects on animals’ incentive evaluation of the instrumental outcome (Parkes et al. 2016), would be an optimal approach to devalue the reinforcer for these experiments. Unfortunately, attempts by our group to establish a reliable sensory-specific satiety devaluation procedure have been unsuccessful (J. S. Innes, unpublished findings; B. J. Everitt, personal communication).

The conclusion that female place learners are more prone to develop S-R instrumental habits than their response learner counterparts would be convergently supported by the demonstration that place learners show a lesser reduction in their instrumental response rate upon the degradation of the instrumental contingency by the provision of response-noncontingent deliveries of the reinforcer (Dickinson and Mulatero 1989; Hammond 1980).

The present findings, interpreted in this way, bear on *1)* the associative nature of place and response navigation strategies and *2)* the relationship between declarative and procedural memory systems and distinct goal-directed and habitual instrumental learning strategies.

There is an extensive literature that utilises response navigation as a tool to study S-R habits, implicitly assuming that response learning is acquired by the strengthening of association between representations of stimulus (e.g., maze choice point) and some motor output (e.g., a left turn; e.g., Ferragud et al. 2010; Packard 1999; Packard and McGaugh 1996; Smith et al. 2012; Smith and Graybiel 2013). This contrasts to place learning, which is variably understood to be acquired through A-O instrumental learning (Kosaki et al. 2018), a model-based learning process (Chersi and Burgess 2015; Geerts et al. 2020), or a non-associative learning mechanism (Cheng 1986; Doeller et al. 2008; Hirsh 1974; O’Keefe and Nadel 1978). Although this widely accepted associative structure for egocentric spatial navigation has historical precedent – whether animals utilise response strategies for navigation was used historically to test S-R reinforcement theories of learning that presently underpin S-R habits (Blodgett and McCutchan 1948; Goodman 2021; Tolman et al. 1946) – it has been rarely tested empirically.

One study has reported that response but not place learning displays some of the defining behavioural features of instrumental habits, such as a resistance to offline changes in the incentive value of the outcome (Kosaki et al. 2018). Here, male rats trained to asymptotic levels of performance in a dual-solution navigation task underwent two “direct opposition” tests, before which they were pre-fed to satiety with either the navigational outcome or an alternative outcome. Sensory-specific satiety increased the proportion of rats relying on a response as opposed to a place learning strategy. It also increased decision latencies in the test, except in the case of rats with hippocampal lesions that could only rely on response learning to solve the discrimination. These results are in line with spatial performance under response but not place navigation strategies being devaluation-resistant (see also Sage and Knowlton 2000).

An issue clouding the interpretation of this kind of evidence is that the exclusively instrumental nature of spatial behaviour (or indeed any exploratory behaviour) is not unambiguous (Dickinson 1994). As the approach of appetitive outcomes is an innate behaviour, there is probably a Pavlovian contribution to learned spatial behaviours difficult to disentangle from instrumental contributions. Indeed, under some conditions learned goal approach behaviours can be remarkably resistant to the introduction of omission contingencies (Hershberger 1986; Williams and Williams 1969), which is expected to disrupt an instrumentally conditioned behaviour (Davis and Bitterman 1971). That some Pavlovian behaviours can resist the devaluation of the US makes it challenging to conclude from the devaluation-resistant nature of spatial behaviours that such behaviours meet the criteria for an instrumental habit.

The approach taken in the present study, based on the assessment of individual differences, is designed to circumvent this particular issue. If response, but not place learning, meets the definition of an S-R instrumental habit, one would expect that those animals tending to rely on response as opposed to place learning strategies for spatial navigation would more readily engage S-R associative control over free-operant lever pressing, a well-established instrumental behaviour. This study did not confirm this prediction, and, in females, found the opposite (**Figure 3E**).

Therefore, the results of this study are more in keeping with the notion that response spatial learning and S-R instrumental habits are dissociable behavioural constructs. This view is supported by studies that have found that the devaluation-resistant properties of spatial behaviour are determined by the amount of training (Smith et al. 2012; Smith and Graybiel 2013; De Leonibus et al. 2011; Sage and Knowlton 2000). In one of these, male rats were trained in a T-maze spatial discrimination task conducted under conditions strongly promoting the reliance on a response spatial learning strategy (Barnes et al. 2011; Smith and Graybiel 2013). Spatial performance was disrupted by conditioning of a taste aversion to the navigational reward after learning criterion was achieved, but not after an additional 10 daily training sessions, even though this additional training did not improve discrimination accuracy.

Neuropsychological studies of patients with brain lesions and controlled lesion studies in non-human animals support the distinction between declarative and procedural types of memory underpinned by distinct neural systems, one instantiation of which is the distinction between place and response navigation strategies (Cohen and Eichenbaum 1993; Cohen and Squire 1980; Milner et al. 1968; Scoville and Milner 1957; Squire 2004; White and McDonald 2002). Similarly, behavioural studies using the free-operant task in rodents have revealed the distinction between goal-directed and habitual instrumental behaviour with distinct associative structures and neural substrates (Balleine and O’Doherty 2010; Dickinson 1985). Given similarities in behavioural features, predictive factors and causative brain regions (see above and *Introduction*), some authors have suggested that declarative memory and goal-directed actions, on one hand, and procedural memory and S-R habits, on the other, are underpinned by the same two-fold systems architecture in the brain (Chersi and Burgess 2015; Geerts et al. 2020). One would predict from this that the tendency to rely on procedural versus declarative memory systems would increase the likelihood to rely on S-R versus A-O control over instrumental behaviour, but this is not supported by the present results (**Figure 3E**).

Indeed, although it is theoretically challenging to account for goal-directed instrumental behaviour without description of some declarative memory process (Dickinson 1980), the declarative memory system does not appear to be essential for goal-directed actions *per se*. Several authors have demonstrated that free-operant lever pressing is equally sensitive to devaluation after extensive lesions to the hippocampal formation, including the entorhinal cortex (Corbit et al. 2002; Corbit and Balleine 2000; Lex and Hauber 2010; Piquet et al. 2024).

Instead, goal-directed instrumental behaviour is in some situations supported by psychological processes dependent on the declarative memory system. For example, the entorhinal cortex and ventral hippocampus are required for goal-directed instrumental responding to adapt to changes in the response-outcome contingency introduced by response-free outcome delivery (Corbit et al. 2002; Corbit and Balleine 2000; Devenport 1979, 1980; Piquet et al. 2024). Further, in instrumental tasks wherein the probability that a response is rewarded is conditional on previous behaviours, goal-directed decision-making requires the chaining of multiple A-O associations together, potentially engaging the relational memory function of the hippocampus (Miller et al. 2017; Vikbladh et al. 2019). This particular fact may explain why individuals relying more on declarative versus procedural spatial learning strategies are more predisposed to engaging model-based instead of model-free reinforcement learning strategies in a two-step probabilistic decision making task (Vikbladh et al. 2019), but are not more likely to rely on A-O-mediated instrumental strategies in the present (one-step) free-operant task (**Figure 3E**).

One can only speculate as to why, in the present experiments, place learners were actually *more* predisposed than response learners to rely on S-R control over instrumental behaviour. Studies in humans suggest that individuals relying more on declarative versus procedural navigation strategies exhibit lower levels of prefrontal cortical-dependent executive functions – for example, working memory capacity and cognitive flexibility (Aumont et al. 2019a, 2019b) – features known to predispose towards the reliance on S-R habits or model-free decision-making under stress (Otto et al. 2013; Quaedflieg et al. 2019). Although rats were not explicitly stressed in the present free-operant task, instrumental conditioning under VI intervals of reinforcement may be stressful to rats by virtue of the unpredictability of reward delivery (Garr et al. 2020), and can also readily entrain adjunctive behaviours such as polydipsic drinking (De Boer et al. 1990; Falk 1961), which are believed to have a stress coping function (Brett and Levine 1979; Marti-Prats et al. 2023). Future research is needed to test this hypothesis.

Further research will be necessary to establish why it is only in females that subjects that tend to rely on declarative versus procedural spatial learning strategies are more prone to engage the habit system in an instrumental setting (**Figure 3E**). Lever pressing was greatly more devaluation-sensitive in males than in females (**Supplementary Figure 2D**), in line with previous studies (Mohammed et al. 2025; Quinn et al. 2007; Schoenberg et al. 2019, 2022), and, as such, instrumental responding may have been “too goal-directed” for any appreciable effect of place/response learning tendency to emerge in males in the present experiments.

Taken together with the existing literature, the results of this study suggest a complex relationship between declarative and procedural memory systems on the one hand, and A-O- and S-R-mediated instrumental strategies on the other, that crucially depend on the specifications of the instrumental task. Thereby, they have profound implications for our understanding of multiple long-term memory systems across different behavioural settings. They also bear on the psychological and neural basis of individual vulnerability to compulsive disorders, in particular how place/response learning tendency may be considered to shape the individual vulnerability to compulsive behaviours (Bohbot et al. 2013; Gregory et al. 2015), which are conceptualised by many as the loss of control over S-R habits (Belin et al. 2013; Everitt et al. 2001; Everitt and Robbins 2005; Gillan et al. 2016; Robbins et al. 2024). Future work should utilise a range of high-quality behavioural models of compulsive disorders to determine the vulnerability to develop compulsive behaviours in animals characterised at baseline for their tendency to rely on declarative versus procedural memory systems, for instance those relying on place versus response strategies in dual-solution navigation tasks.

## Methods and materials

### Subjects

Seventy-two male and female Sprague Dawley rats (36 females; *Charles River, UK*), weighing between 315-425 g (males) and 235-335 g (females) by the start of experiments, were housed in groups of three or four under a reversed 12-hour light/dark cycle (lights off at 7 AM) in a temperature-(21 ± 1 °C) and humidity-controlled (∼55%) environment. After seven days of habituation to the animal facility, rats were single-housed and food-restricted to gradually reach ∼85% of their predicted free-feeding body weight before starting behavioural training, a level of food restriction maintained throughout the experiments. Water was available *ad libitum* in the home cage. Experiments were performed seven days per week between 8AM and 7PM. All experimental protocols were conducted under Home Office project licence PP0463796 held by Professor David Belin in accordance with the regulatory requirement of the UK Animals (Scientific Procedures) Act (ASPA) 1986, amendment regulations 2012, following ethical review by the University of Cambridge Animal Welfare and Ethical Review Body (AWERB).

### Timeline of the experiments

The experiments conducted in this study are schematically summarised in **Figure 1A**.

### Dual-solution T-maze task

#### Apparatus

The dual-solution T-maze task was conducted in a dedicated, large trapezoidal room (**Supplementary Figure 3**; a = 200 cm, b = 320 cm, h = 290 cm) in bright light. The T-maze (*Viewpoint Life Science, France*) was built from black, infrared-translucent Plexiglas and configured into three enclosed arms of equal dimensions (height = 30 cm; width = 10 cm; length = 45 cm) arranged into the shape of a “T”: a starting arm, and, at right angles, two lateral arms. Slots for guillotine doors enabled the experimenter to block access to each lateral arm.

The T-maze was positioned onto a white, Plexiglas infrared floor (110 cm^2^; *Viewpoint Life Science, France*) containing an array of infrared light-emitting diodes that backlit the maze and the animal. Infrared light was detected by a ceiling-mounted camera fitted with an infrared filter (*Viewpoint Life Science, France*), itself connected to a computer in an adjacent control room.

Commercial software (*Viewpoint Life Science, France*) was used to time the behavioural procedures and live-track the experimental subjects. The software used a pixel threshold approach, which made use of the infrared opacity of animals’ bodies in relation to the T-maze. Frames were pixel-thresholded according to a user-defined cut-off, and the remaining “black” pixels averaged in space to define the “centre” of the animal. The regions of interest manually specified to the software corresponded to each arm and the central intersection of the T-maze.

Pilot experiments using the dual-solution T-maze task showed that a large majority of rats adopted a response strategy. To enhance the heterogeneity of visual stimulation in the extra-maze environment, increasing the likelihood that a subject would rely on place learning (Packard and Goodman 2013; Restle 1957), four laminated cards (38 x 45 cm) displaying different black-and-white patterns were positioned slightly outside of, and parallel to, each side of the infrared platform, roughly 2 m above the ground (**Supplementary Figure 3**). This was achieved by sticking cards using blue tac onto custom-made, wooden stands. Such cues were never placed in line with a lateral arm so to discourage the acquisition of the spatial discrimination by a cue learning strategy towards an extra-maze cue (O’Keefe and Nadel 1978; Packard and McGaugh 1996; Tolman et al. 1946).

#### Habituation

To reduce the neophobic response to the task reward, rats were firstly habituated to the test pellets used in the dual-solution T-maze task (*5TUL 45mg tablet #1811155, TestDiet, USA*), with 20 pellets delivered to the home cage 24 hours before exposure to the T-maze. Rats were subsequently habituated to the maze in three daily sessions. Firstly, they were placed in the centre of the maze and left to explore its entirety for 15 minutes with ten pellets at the end of each arm. The next day, rats were placed in the starting arm for three five-minute phases: for the first, one of the lateral arms (counterbalanced) was closed off from the starting arm by a guillotine door, and the other contained ten pellets at its end; for the second, this initial configuration was reversed; finally, both lateral arms containing 10 pellets were made accessible. Finally, rats were placed in the starting arm and left to explore the full T-maze for 15 minutes with ten pellets at the end of each lateral arm. For these habituation sessions, the experimenter was outside the room. The maze was cleaned with 10% ethanol between rats.

The third habituation session was employed to assign each individual to the lateral arm, left or right, that would be baited in training. Lateral arm preference was calculated as follows.

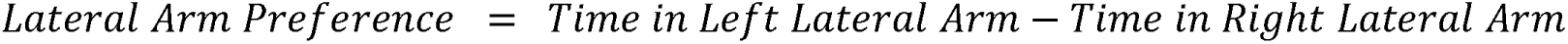

In descending order of preference, males and females separately were assigned their non-preferred arm until half of each sex were assigned to one specific lateral arm. Then they were assigned their preferred arm to counterbalance baited lateral arm assignment within sexes.

#### Training

Rats were trained over at least two daily training sessions. Before each session, two pellets were placed at the end of the rat’s assigned baited lateral arm. The rat was brought into the room and left in a carrying box with the lid removed in front of and roughly at the same height as the floor of the starting arm of the T-maze for two minutes. To start a trial, the rat was placed at the end of the starting arm, with both lateral arms accessible. The experimenter scored the trial from outside the room, watching a computer monitor. Trials were scored as follows.

1) A *correct trial* if the rat entered the baited lateral arm directly from the starting arm.
2) An *incorrect trial* if the rat entered the non-baited lateral arm directly from the starting arm but tracked back to eventually enter the baited lateral arm.
3) An *omission trial* if the rat remained in the starting arm for the entire two-minute trial. The rat was judged to have entered a lateral arm if the entirety of its body, excluding its tail, passed laterally to an imaginary line that projected from each edge of the starting arm. Regardless of how it was scored, each trial was terminated as soon as the rat entered the baited lateral arm, or otherwise after two minutes. With a 10-second delay to allow it to consume the pellets, the rat was removed to the carrying box for a 30-second inter-trial interval (ITI), during which the maze was cleaned with 10% ethanol and the two pellets replaced for the next trial. Rats that had not consumed the pellets were guided by the experimenter to the end of the baited lateral arm, where they could consume the pellets, once the trial ended.

All subjects received two daily sessions of six trials in the dual-solution T-maze task. Rats that discriminated to a learning criterion of five consecutive correct trials within a session received no further training (Ferragud et al. 2010). The remaining subjects received an additional four trials on the second day of training, along with two daily sessions of up to 15 trials. This schedule concluded as soon as rats achieved the learning criterion (Ferragud et al. 2010).

#### “Direct opposition” testing

To determine the learning strategy, place or response, on which rats relied to solve the dual-solution spatial discrimination, a “direct opposition” test was conducted under extinction conditions with the T-maze rotated by 180° (**Figure 1**; Blodgett and McCutchan 1948; Packard and McGaugh 1996; Restle 1957). Rats were tested in a single trial occurring 24 hours after attaining learning criterion in training (see above). The rat was brought into the room and left in the carrying box for two minutes in the same location as in training. To start the test, the rat was positioned at the end of the starting arm, with both lateral arms accessible. Watching a monitor outside of the room, the experimenter judged which lateral arm the rat entered first.

As illustrated in **Figure 1B**, rats were judged to have relied on a place strategy to solve the dual-solution T-maze task if they selected the lateral arm in the same allocentric location, as defined in relation to the geometry of the room and the extra-maze cues, as the baited arm in training. This lateral arm was in the opposite egocentric relationship to the starting arm as the baited lateral arm, i.e., place learners turned left when they were trained to turn right, or *vice versa*. Rats that selected the lateral arm in the same egocentric relationship to the starting arm as the baited lateral arm in training were judged to have relied on response learning.

### Instrumental food task

#### Apparatus

Instrumental food task training and testing occurred in 24 dedicated extra-tall operant chambers (53.3 cm x 34.9 cm x 27.3 cm; *Med Associates, USA*) located within ventilated sound-attenuating cupboards. Operant chambers were outfitted with a house light, a food magazine through which dustless food pellets (*5TUL 45mg*) could be delivered, and a single retractable metal lever flanking the magazine to the left or right (counterbalanced). Infrared beams were installed in the magazines to record magazine head entries. Chambers were controlled by *MedPC IV* software (*Med Associates, USA*).

#### Training

Rats were habituated to the operant chambers with a 30-minute session signalled by the house light and the next day trained to retrieve the test pellets (*5TUL 45mg*) from the food magazine in a single session of 30 deliveries under a variable time (VT) 30s schedule, with no *operanda* available in either session. Subsequently, rats were trained to press the lever to receive a pellet under a fixed ratio (FR) 1 schedule of reinforcement in the first two sessions, a variable interval (VI) 20s schedule in the next two sessions, and a VI 60s schedule in the remaining five (*Experiment 1*) or 20 (*Experiment 2*) sessions. Lean VI schedules are conducive to the development of S-R habits (DeRusso 2010; Dickinson et al. 1983). A training session ended after 30 pellet deliveries or a time limit of 30 minutes (FR1, VI 20s) or 60 minutes (VI 60s).

#### Devaluation testing

To assess the degree to which instrumentally conditioned behaviour was goal-directed versus habitual, the sensitivity of lever pressing to the pre-feeding of the reinforcer to satiety was measured in a within-subjects design (Colwill and Rescorla 1985; Gourley et al. 2010).

Starting roughly 22 hours after subjects’ final VI 60s conditioning session, rats underwent a four-day devaluation test. The cohort was divided into two equally sized groups of males and females. On the first day of the test, the first group received *ad libitum* access to the test pellets (≥60 g in a metallic cup) in the home cages (the devalued condition). The second group was left undisturbed (the valued condition). After two hours, rats were removed from their home cages and underwent a five-minute test conducted in the operant chambers under extinction conditions; that is, the lever was available and operable but had no programmed consequences. After two daily VI 60s sessions to re-baseline instrumental performance, the first day of the devaluation test was repeated once; here, the second group was pre-fed.

To determine how effective pre-feeding to satiety was at reducing the value of the reinforcer in *Experiment 2*, a consumption test was conducted immediately after the instrumental response rate was assayed. All rats, irrespective of group, received *ad libitum* access to 70 g of the test pellets in a food cup in the operant boxes for a 15-minute period signalled by the illumination of the house light. Cups were weighed before and after to calculate reinforcer consumption.

### Exclusions

Seven subjects (two females) in *Experiment 1* and three subjects (one female) in *Experiment 2* failed to attain the learning criterion in the dual-solution T-maze task and were consequently excluded from all analyses. In *Experiment 2*, one male subject that contributed an outlier data point in the devaluation test (the number of lever presses in the first minute under the devalued condition: 3.52 standard deviations greater than the mean) was excluded from all analyses.

## Data and statistical analyses

### Data were analysed using custom-written code in R

When required, assumptions for normal distribution, homoscedasticity and sphericity were verified using the Shapiro-Wilk, Levene, and Mauchly sphericity tests, respectively. For some analyses, reliable estimation of sphericity using Mauchly’s test required the aggregation of time series data into blocks to reduce the number of levels of the within-subjects factor. When the mentioned assumptions were violated, data were transformed using the square root function. In such cases, data are presented as they were transformed throughout the manuscript.

Figures were created using custom-written code in *matplotlib* (Hunter 2007) and *seaborn* (Waskom 2021) packages in Python. Data are presented in figures as means ± standard error of the mean (SEM) or box plots (medians ± 25% percentiles as boxes, range as whiskers).

The significance level was set at α = 0.05.

Effect sizes are reported as partial eta squared (η_p_ ^2^) or odds ratio (OR).

As they involved different subjects, *Experiments 1* and *2* were analysed separately.

### Dual-solution T-maze task performance

The number of trials required for subjects to attain the learning criterion was analysed using factorial ANOVAs. As video recording was unavailable, there was no latency data for this task.

The distribution of place and response learners in male and female subjects was analysed using chi-squared (χ^2^) contingency tests. When the expected frequency rule (≥ 20% of cells have a frequency of ≥ 5; Cochran 1954) was violated, Fisher exact tests were instead performed.

### Instrumental training performance

The rate of lever presses and magazine entries across instrumental training sessions were analysed using mixed ANOVAs. In *Experiment 2*, data points were amalgamated into two-session blocks to reduce the levels of the within-subjects factor to ensure a satisfactory reliability of the estimation of sphericity using Mauchly’s test. Statistically significant interactions were followed up with *post hoc* tests with Tukey’s correction where appropriate. *Devaluation test performance*

The number of lever presses and magazine entries during the first minute of the five-minute test across devalued and valued conditions were analysed using mixed ANOVAs. Significant interactions were followed up with *post hoc* tests with Tukey’s correction where appropriate. Analyses were restricted to the first minute of the five-minute test as pilot experiments demonstrated a pronounced decline in the rate of instrumental responding across the course of the test, probably reflecting instrumental extinction learning (M. Fouyssac, unpublished findings; J. S. Innes, unpublished findings). It was desired to isolate the effects of reinforcer devaluation from those of instrumental extinction learning on the instrumental response rate. *Consumption test performance*

The free consumption of the reinforcer during the consumption test across devalued and valued conditions were analysed using mixed ANOVAs. Statistically significant interactions were followed up using *post hoc* tests, correcting using Tukey’s method where appropriate.

## Acknowledgements

This work, carried out at the Department of Psychology and the Department of Physiology, Development and Neuroscience of the University of Cambridge, was supported by grants from UKRI-Medical Research Council (MR/W019647/1) and the Rosetrees Trust (OoR2023\100003) to DB. JSI was supported by a grant from the Rosetrees Trust (MB2022\100002). The authors would like to thank Mickaël Puaud for his technical assistance.

## Authorship Contributions

DB and JSI designed the experiments. JSI carried out the experiments and formally analysed the data. JSI wrote the manuscript with critical input from DB.

## Data availability

Data will be made available on the University of Cambridge repository after acceptance of the manuscript for publication.

