## Supplementary Information for "Relying on a declarative as opposed to a procedural learning strategy predisposes to the development of instrumental habits in female but not in male rats"

**Supplementary Figure 1**

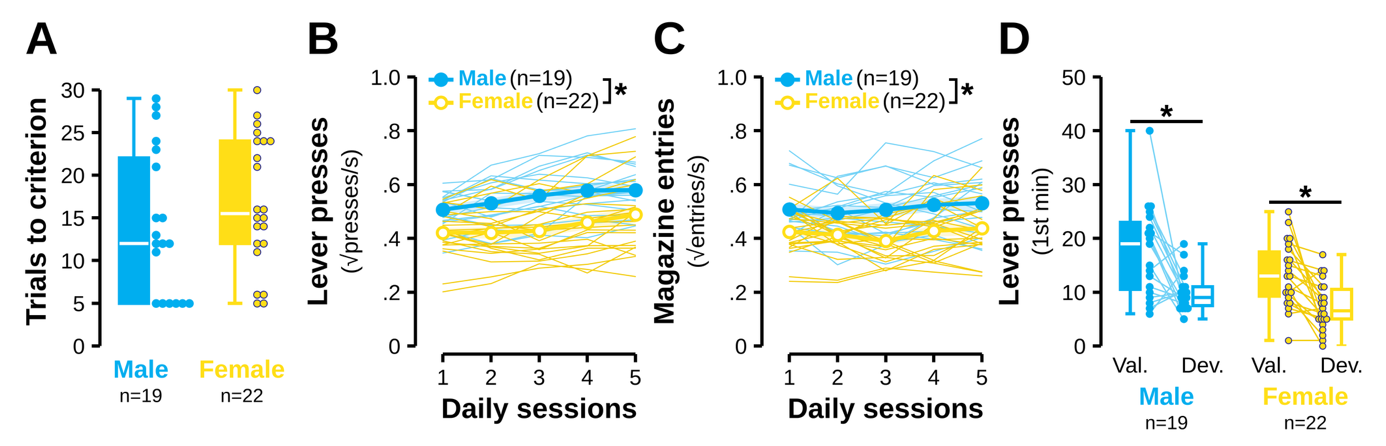

**Supplementary Figure 1: After a limited period of training, the instrumental response is more frequent but equally resistant to reinforcer devaluation in males and females.**

**A)** Males and females acquired the T-maze discrimination at the same rate, requiring an equivalent number of trials to attain the learning criterion (for all F, F < 1). **B-C)** Across five daily sessions of instrumental conditioning under a variable interval (VI) 60s schedule of reinforcement, males pressed the lever (*B*; Sex: F_1,37_ = 8.07, p = 0.007, η_p_^2^ = 0.18; Sex x Day: F_2.29,84.73_ = 1.90, p = 0.15, η_p_^2^ = 0.049) and entered the food magazine (*C*; Sex: F_1,37_ = 7.98, p = 0.008, η_p_^2^ = 0.18; Day: F_2.70,99.72_ = 1.84, p = 0.15, η_p_^2^ = 0.047; Sex x Day: F_2.70,99.72_ < 1) at a greater frequency than their female counterparts. **D)** In a post-training devaluation test, the pre-feeding to satiety of the reinforcer (the devalued condition; Dev.) reduced the instrumental response rate during the first minute of a five-minute test conducted under extinction conditions compared to no pre-feeding (the valued condition; Val.; Condition (min1): F_1,37_ = 21.99, p < 0.001, η_p_^2^ = 0.37). This effect was equal across sexes (Sex x Condition (min1): F_1,37_ < 1).

**Supplementary Figure 2**
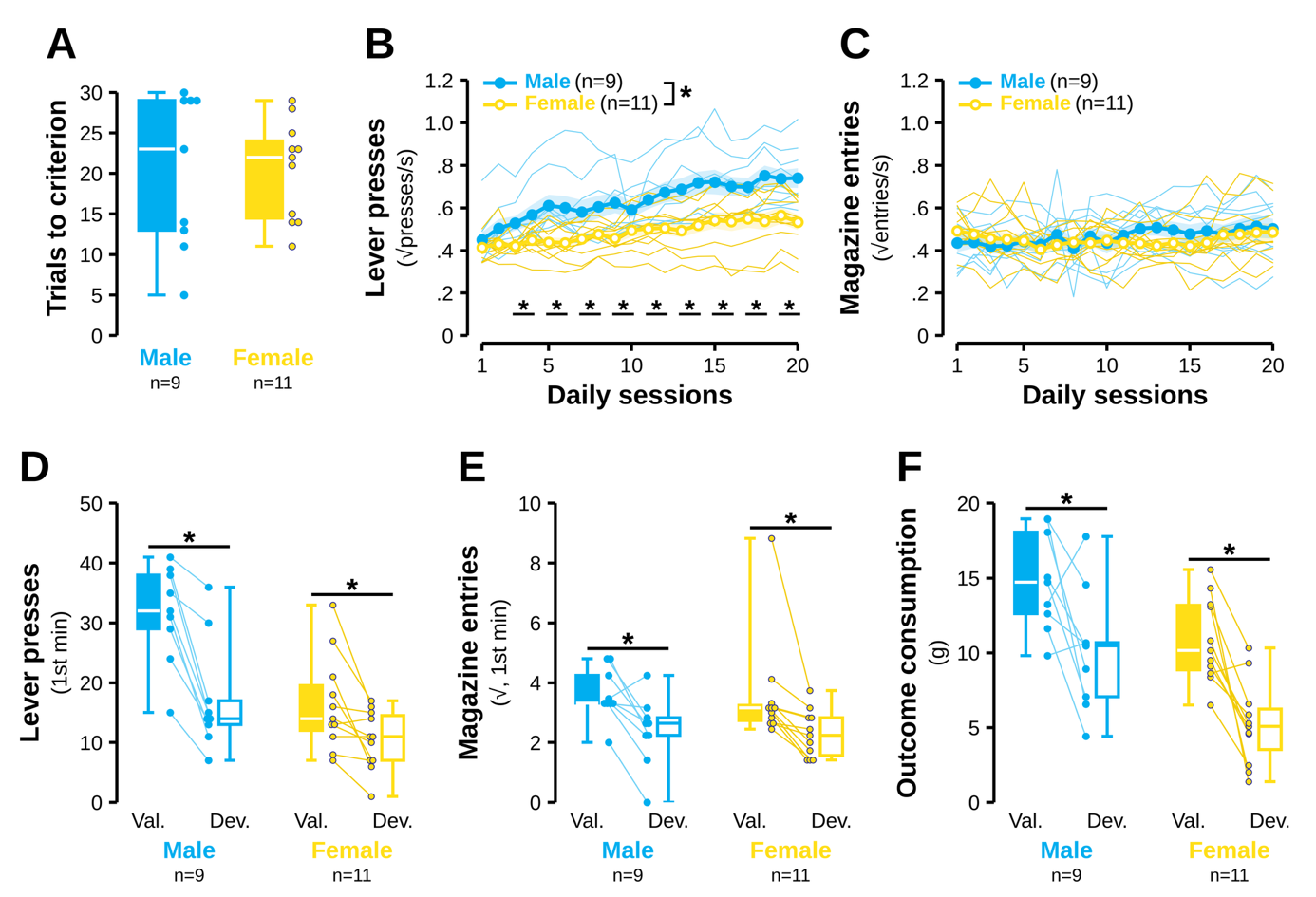

**Supplementary Figure 2: Instrumental behaviour is more resistant to reinforcer devaluation in females than in males after extended instrumental conditioning.**

**A)** Males and females acquired the T-maze discrimination at the same rate, requiring an equivalent number of trials to attain the learning criterion (for all F, F < 1). **B)** Overall, across 20 daily sessions of instrumental conditioning under a variable interval (VI) schedule of reinforcement, the cohort increased their lever pressing rates (Two-Day Block: F_2.72,43.49_ = 33.83, p < 0.001, η_p_^2^ = 0.68). This increase was profoundly greater in male rats than in their female counterparts (Sex: F_1,16_ = 10.49, p = 0.005, η_p_^2^ = 0.40; Sex x Two-Day Block: F_2.72,43.49_ = 4.02, p = 0.016, η_p_^2^ = 0.20): both sexes pressed the lever with equal frequency over the first two training sessions (p = 0.14); but, by the final two sessions, the instrumental response rate was substantially greater in males than in females (p = 0.002). **C)** Males and females entered the food magazine at the same frequency throughout training (Sex: F_1,16_ < 1; Sex x Two-Day Block: F_3.75,60.06_ = 1.65, p = 0.18, η_p_^2^ = 0.094). **D)** In a post-training devaluation test, the pre-feeding to satiety of the reinforcer (the devalued condition; Dev.) reduced the instrumental response rate during the first minute of a five-minute test conducted under extinction conditions compared to no pre-feeding (the valued condition; Val.; Condition (min1): F_1,16_ = 51.40, p < 0.001, η_p_^2^ = 0.76). Although the rate of instrumental responding was attenuated by reinforcer devaluation in both sexes, the effect was less pronounced in females than in males (Sex x Condition (min1): F_1,16_ = 12.01, p = 0.003, η_p_^2^ = 0.43; *post hoc* (males): p < 0.001; *post hoc* (females): p = 0.008). **E)** Non-instrumental behaviours such as magazine entries were also attenuated by the devaluation of the instrumental outcome (Condition (min1): F_1,16_ = 20.40, p < 0.001, η_p_^2^ = 0.56). However, magazine entries were equally sensitive to reinforcer devaluation in males and females (Sex x Condition (min1): F_1,16_ < 1). **F)** As expected, rats freely consumed a smaller quantity of the reinforcer after it was pre-fed to satiety than after no intervention, this being measured across a 15-minute assay starting immediately after the instrumental response rate was assessed in the devaluation test (Condition: F_1,16_ = 24.87, p < 0.001, η_p_^2^ = 0.61). The effect of reinforcer devaluation on the free consumption of the instrumental outcome did not differ between males and females (Sex x Condition: F_1,16_ < 1).

**Supplementary Figure 3**

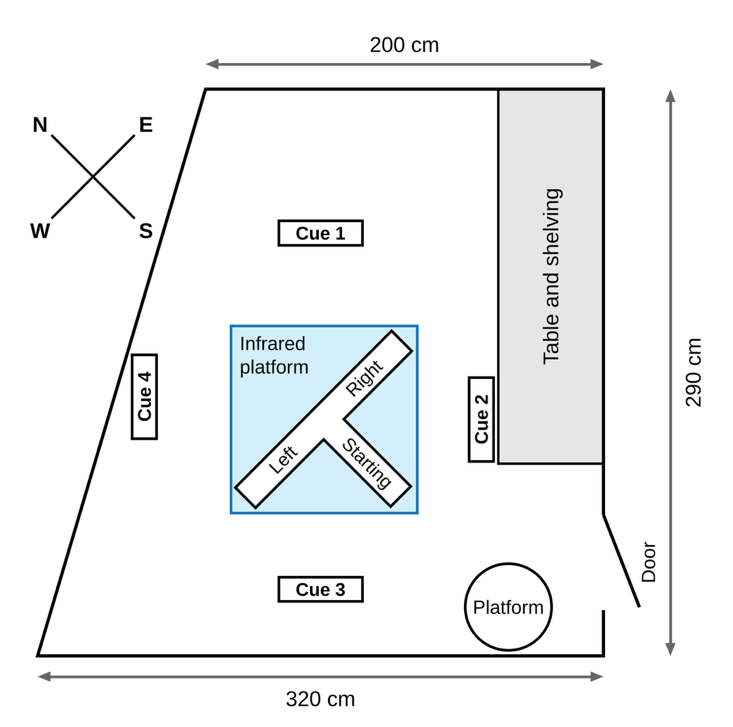

**Supplementary Figure 3: Layout of the navigational environment for the dual-solution T-maze task.** The dual-solution T-maze task was carried out in a dedicated video-tracking room with a trapezoid geometry (a = 200 cm, b = 320 cm, h = 290 cm). The T-maze (height = 30 cm; arm width = 10 cm; arm length = 45 cm) was positioned roughly in the centre of the room on an infrared platform (110 cm^2^). Extra-maze visual cues included four laminated cards (38 cm x 45 cm), which were placed in a square pattern surrounding the T-maze roughly 2 m above the ground. This was achieved by sticking cards onto custom-made, wooden stands. Prominent shelving and a table lined the side of the wall next to the door. To start a training session or test trial, the rat was brought into the video tracking room in a carrying box and placed on a platform in front of the starting arm of the maze. This is also where the rat resided during the inter-trial interval, during the maze was cleaned and the food reward replaced.
